# A thermal trade-off between viral production and degradation drives phytoplankton-virus population dynamics

**DOI:** 10.1101/2020.08.18.256156

**Authors:** David Demory, Joshua S. Weitz, Anne-Claire Baudoux, Suzanne Touzeau, Natalie Simon, Sophie Rabouille, Antoine Sciandra, Olivier Bernard

**Author notes:** Contributed equally (Joshua S. Weitz, Anne-Claire Baudoux, Suzanne Touzeau).

## Abstract

Marine viruses interact with their microbial hosts in dynamic environments shaped by variations in abiotic factors, including temperature. However, the impacts of temperature on viral infection of phytoplankton are not well understood. Here we coupled mathematical modeling with experimental datasets to explore the effect of temperature on three *Micromonas*-prasinovirus pairs. Our model shows the negative consequences of high temperatures on infection and suggests a temperature-dependent threshold between viral production and degradation. Modeling long-term dynamics in environments with different average temperatures revealed the potential for long-term host-virus coexistence, epidemic free, or habitat loss states. Hence, we generalized our model to global sea surface temperature of present and future seas and show that climate change may influence virus-host dynamics differently depending on the virus-host pair. Our study suggests that temperature-dependent changes in the infectivity of virus particles may lead to shifts in virus-host habitats in warmer oceans, analogous to projected changes in the habitats of macro- and micro-organisms.

## Introduction

Viruses are the most abundant biological entity in the ocean (Suttle 2005, 2007). Viruses are present in all ocean environments, even in the most extreme biomes (Williamson *et al*. 2008), and play essential roles in ecosystem processes (Breitbart *et al*. 2018, Fuhrman 1999, Jover *et al*. 2014, Wilhelm & Suttle 2000). For example, viruses of phytoplankton are responsible for significant mortality of their hosts in coastal and open ocean (Baudoux *et al*. 2007, Bratbak *et al*. 1998) and enhance microbial diversity (Thingstad 2000, Weinbauer & Rassoulzadegan 2004). They indirectly contribute to nutrient recycling (Jover *et al*. 2014) through the release of dissolved organic matter upon host lysis. Despite this substantial impact on phytoplankton communities, viral infection dynamics is far from being understood (Knowles *et al*. 2016).

Abiotic factors such as temperature, salinity, nutrients and light can affect viral infection processes that drive phytoplankton-virus dynamics (Mojica & Brussaard 2014). These environmental factors impact infection processes at different levels, ranging from host metabolism, viral structure, viral infectivity to viral life strategy (Baudoux & Brussaard 2005, Cordova *et al*. 2003, Tomaru *et al*. 2005, Weinbauer *et al*. 2003, 1999, Wigginton *et al*. 2012, Wilhelm *et al*. 1998). However, relatively few studies have explicitly considered the impact of temperature on viral dynamics (Mojica & Brussaard 2014).

Previous work suggests that temperature may alter interactions between viruses and their phytoplankton hosts. Elevated temperatures may induce a loss of infectivity among viral particles (Baudoux & Brussaard 2005, Demory *et al*. 2017, Martínez *et al*. 2015, Nagasaki & Yamaguchi 1998), and may also impact phytoplankton metabolism (Ras *et al*. 2013) and resistance to infection (Kendrick *et al*. 2014, Tomaru *et al*. 2014). Field studies have found latitudinal variations of virus-induced mortality in marine microbial assemblages. For example, increasing viral lysis rates of phytoplankton were recorded from high to low latitudes across the North Atlantic Ocean and, interestingly, correlated positively with temperature (Mojica *et al*. 2016). However, it remains unclear whether temperature has a mechanistic role in such differences or whether variation in lysis rates are attributable to other factors, e.g., gradients in viral life history strategies (Brum *et al*. 2016) or variations in microbial community composition (Baudoux *et al*. 2015, Martin-Platero *et al*. 2018).

Understanding the role of temperature on phytoplankton viral infection is essential, especially to predict how virus-host dynamics may change in a warming ocean. To do so, we turned to mathematical models of viruses, microbial hosts and their environment (Weitz 2016). The bulk of modeling work on virus-microbe dynamics focused on phage-bacteria dynamics (Abedon 2008, Jacquet *et al*. 2010, Weitz 2016). Mathematical models often assumed that bacterial populations change due to infection and resource limitations. Dynamical models sometimes included variation in life-history traits, such as burst size and latent period (Middelboe 2000, Middelboe *et al*. 2001), as well as resistance mechanisms (Bohannan & Lenski 1997a). Few biogeochemical models described phytoplankton infection by viruses to explore the extent of viral-mediated nutrient cycling (Béchette *et al*. 2013, Bohannan & Lenski 1997b, Gerla *et al*. 2013, Samanta *et al*. 2013). Other models investigated the role of viruses on ecosystem processes in multitrophic models (Rhodes & Martin 2010, Rhodes *et al*. 2008, Weitz 2016).

Here, we investigated the impact of temperature on the dynamics of *Micromonas*, a widely distributed and abundant picophytoplankton, and their virus by combining a non-linear population model with experimental data. We first accounted for infectious and non-infectious viruses and the impact of temperature on various infection processes. We then parameterized this model with laboratory data for three pairs of species of the picoeukaryote *Micromonas* genus and prasinoviruses: RCC829 (*M. bravo*)/RCC4265 (Mic-B/MicV-B), RCC451 (*M. commoda*)/RCC4253 (Mic-A/MicV-A) and RCC834 (*M. pusilla*)/RCC4229 (Mic-C/MicV-C). Using model-data integration methods, we examined how temperature affects near-term infection dynamics. We then analyzed the long-term dynamics to understand the impact of temperature variability. Finally, we generalized our model to include projected changes in global sea surface temperature (SST) for present and future periods. In doing so, we explored how incorporating temperature into infection processes alters projections of both virus and microbial population dynamics in a changing ocean.

## Results

### Modeling *Micromonas*-virus population dynamics

The model represents phytoplankton infection by a single virus species (Fig. 1). We assumed that phytoplankton growth without viruses can be represented as a logistic function with a carrying capacity *K* and a gross growth rate *μ*. Phytoplankton hosts are subdivided into two sub-populations: susceptible to be infected (*S*) and infected phytoplankton (*I*) so that total host concentration is *N* = *S* + *I*. Susceptible cells are infected by free-living infectious viruses (*V_i_*) at an infection rate *ϕ*. Infected cells are lysed at a rate λ. Both susceptible and infected cells die at a rate *ψ*, due to processes other than viral lysis. The following system of ordinary differential equations (ODE) describes the resulting phytoplankton dynamics:

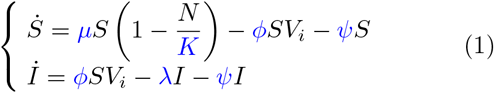

**FIG. 1:**
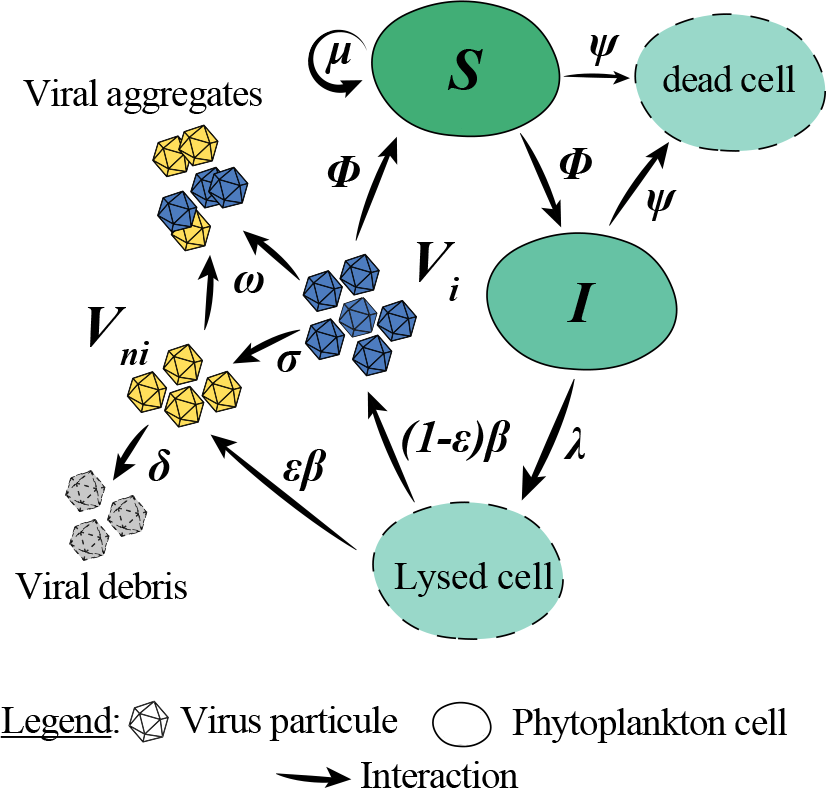
Schematic representation of the model. Susceptible hosts *S* grow at a rate *μ* and become infected at a rate *ϕ*. Infected hosts *I* are lysed at a rate λ and produce viruses in the media with a burst size *β*. Both *S* and *I* die at a rate *ψ*. Viruses are produced in two stages: a fraction (1 – *ϵ*) is infectious *V_i_* and a fraction *ϵ* is non-infectious *V_ni_*. *V_i_* lose their infectivity at a rate *σ* and *V_ni_* are degraded at a rate *δ*. *V_i_* and *V_ni_* particles aggregate at a rate *ω*.

In line with Perelson (2002), we classified the virus into two stages: infectious (*V_i_*) and non-infectious (*V_ni_*). The total number of viruses in their free-living stage in the medium is equal to *V* = *V_i_* + *V_ni_*, and the following ODE system describes their dynamics:

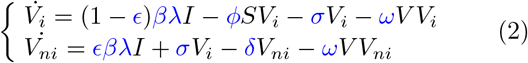

Lysed infected cells produce free viruses at a rate *β*λ. The fraction of infectious viruses produced is (1 – *ϵ*), whereas it is e for the non-infectious viruses. Infectious viruses become non-infectious at a rate *σ* before degrading. Non-infectious viruses are degraded at a rate *δ*. Viral particles can irreversibly aggregate at a rate *ω* due to diffusion and biophysical properties. Supplementary Tab. 1 summarizes model variables and parameters (termed life-history traits). In this “basal” model, life-history traits are constant with temperature.

To describe the impact of temperature on *Micromonas*-virus infection, we extended the basal model by considering each life-history trait as a function of temperature (termed the “temperature-driven model”; see Methods for complete description).

### Virus-host dynamics reproduced by temperature-driven model

To fit our temperature-driven model to the experimental data, we estimated the hyper-parameters of the temperature-driven function using a non-linear fitting algorithm (see Methods). The temperature-driven model was efficiently able to reproduce the experimental data from Demory *et al*. (2017) with the host-virus Mic-B/MicV-B pair (Fig. 2). For the six temperatures tested, we obtained an accurate representation of the host and virus dynamics with lower Akaike information criterion (AIC) and Bayesian information criterion (BIC) for the majority of the temperatures compared to the basal model (Supplementary Tab. 2). We re-calibrated the model for two other Host-Virus pairs: Mic-A/MicV-A and Mic-C/MicV-C. While these new data-sets were not as accurate, the temperature-driven model was able to adequately reproduce the dynamics for five temperatures (Supplementary Fig. 1 and Supplementary Fig. 2).

**FIG. 2:**
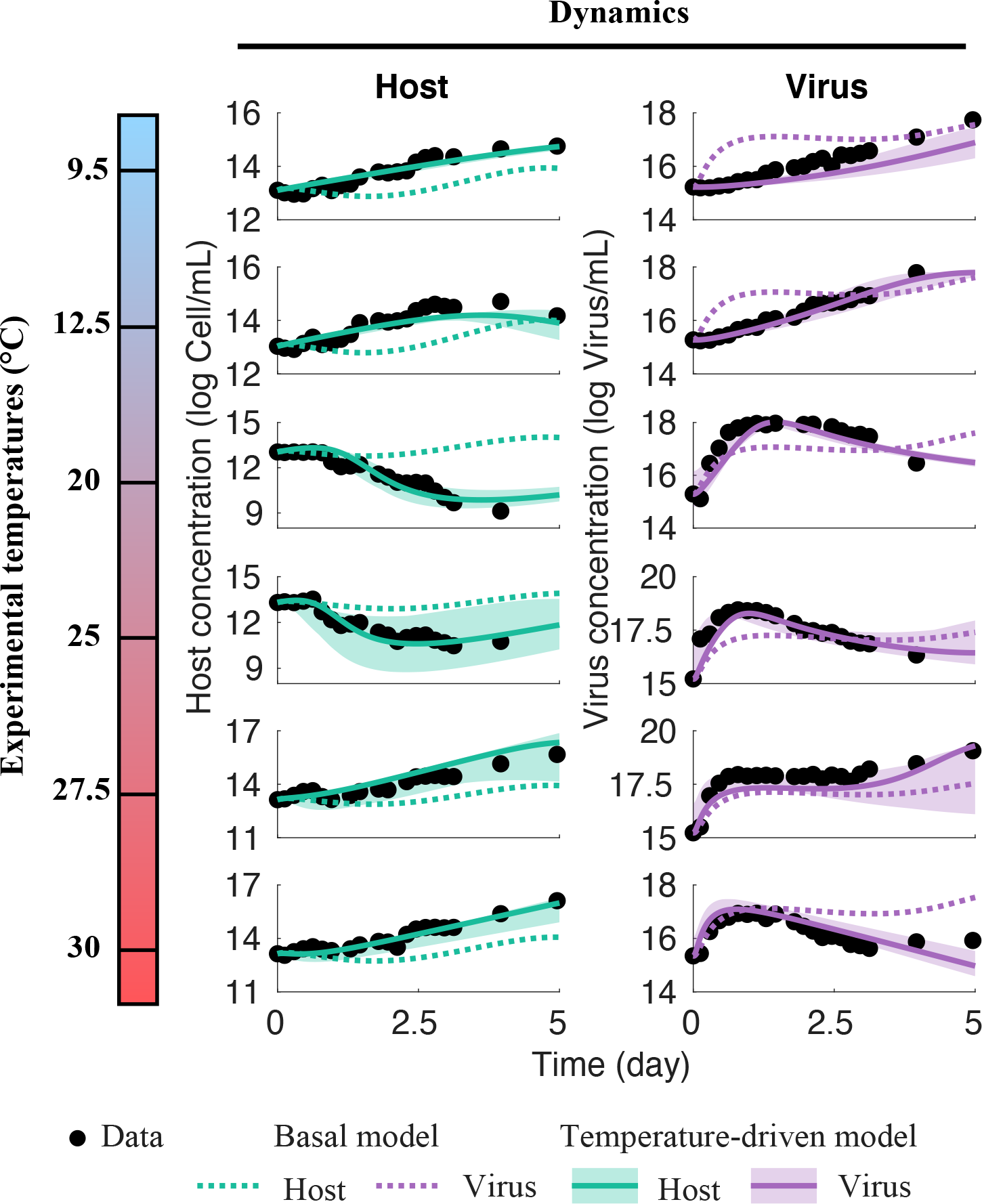
Model fits for the six temperatures tested experimentally in Demory *et al*. (2017). The left column shows the phytoplankton dynamics of the total host population *N* simulated by the model (green line) and the corresponding experimental data (black circles). The right column shows the virus dynamics of the total virus population *V* simulated by the model (purple line) and the corresponding experimental data (black circles). The solid lines are the temperature-driven model with shaded area representing the model fits variability when accounting for a ± 1°C uncertainty on the experimental temperatures. Dashed lines are the basal model.

### Mechanistic inference: validating estimated life history traits

To validate predicted life-history traits, we compared the estimated temperature-driven life-history trait for the pair Mic-B/MicV-B with experimental measurements from Demory *et al*. (2017) and Demory *et al*. (2018) (Fig. 3). When available, the measurements validated our life-history trait estimations for different temperatures. The thermal response of *Micromonas* strain (Fig. 3a) reflected its thermotype (*Micromonas bravo* Warm thermotype), and its thermal environment at isolation site (Naples Bay: 40.75° N, 14.33° E). The optimal temperature was found to be 25.3°C with *μ_opt_* = 0.94 d^−1^, and the maximum growth temperature was 33.5°C. The estimated cardinal temperature 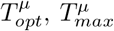 and the *μ_opt_* corresponded to our previous study (Demory *et al*. 2017). Only the estimated 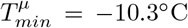 differed from the one calculated in Demory *et al*. (2018) at low temperatures. The estimated latent period and burst size were also well validated by measurements at different temperatures (Fig. 3b and c). We found an optimal temperature for viral lysis close to the optimal host growth temperature 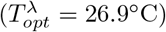, with an optimal lysis rate of about 11.5 d^−1^ (latent period = 2.09 hours) associated with a burst size of 195 viruses released per lysed cell. Below this optimum, the lysis rate increased with temperature, whereas it decreased abruptly beyond. Regarding the percentage of non-infectious produced viruses (*ϵ*, Fig. 3d), we found the half-saturation was equal to 24.7°C < 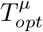, indicating that below the optimal temperature of growth, most viruses produced are infectious. Beyond 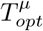, *ϵ* abruptly reaches a plateau where most of the viruses produced are not infectious. Our loss of infectivity (*σ*) and viral decay (*δ*) equations overestimated the experimental rates obtained from data in Demory *et al*. (2017) (by a factor 1 and 0.7 times on average respectively), but represented the exponential qualitative increase with temperature well (Fig. 3e). Estimated adsorption rate (Fig. 3f) increased from 8.6 10^−9^ ml cell^−1^ day^−1^ at 7.5°C to 2.6 10^−7^ ml cell^−1^ day^−1^ at 30°C. Our model adequately captured the increase in adsorption with temperature in a manner similar to biophysical models (Murray & Jackson 1992, Talmy *et al*. 2019). Our adsorption rate estimations were lower than the maximal adsorption rates at all temperatures, suggesting that other biophysical drivers impact adsorption Talmy *et al*. (2019)(Supplementary Fig. 3). The temperature-driven life-history trait for pairs Mic-A/MicV-A, and Mic-A/MicV-A were similar to Mic-B/MicV-B but shifted depending on the host thermal niche (Supplementary Fig. 4).

**FIG. 3:**
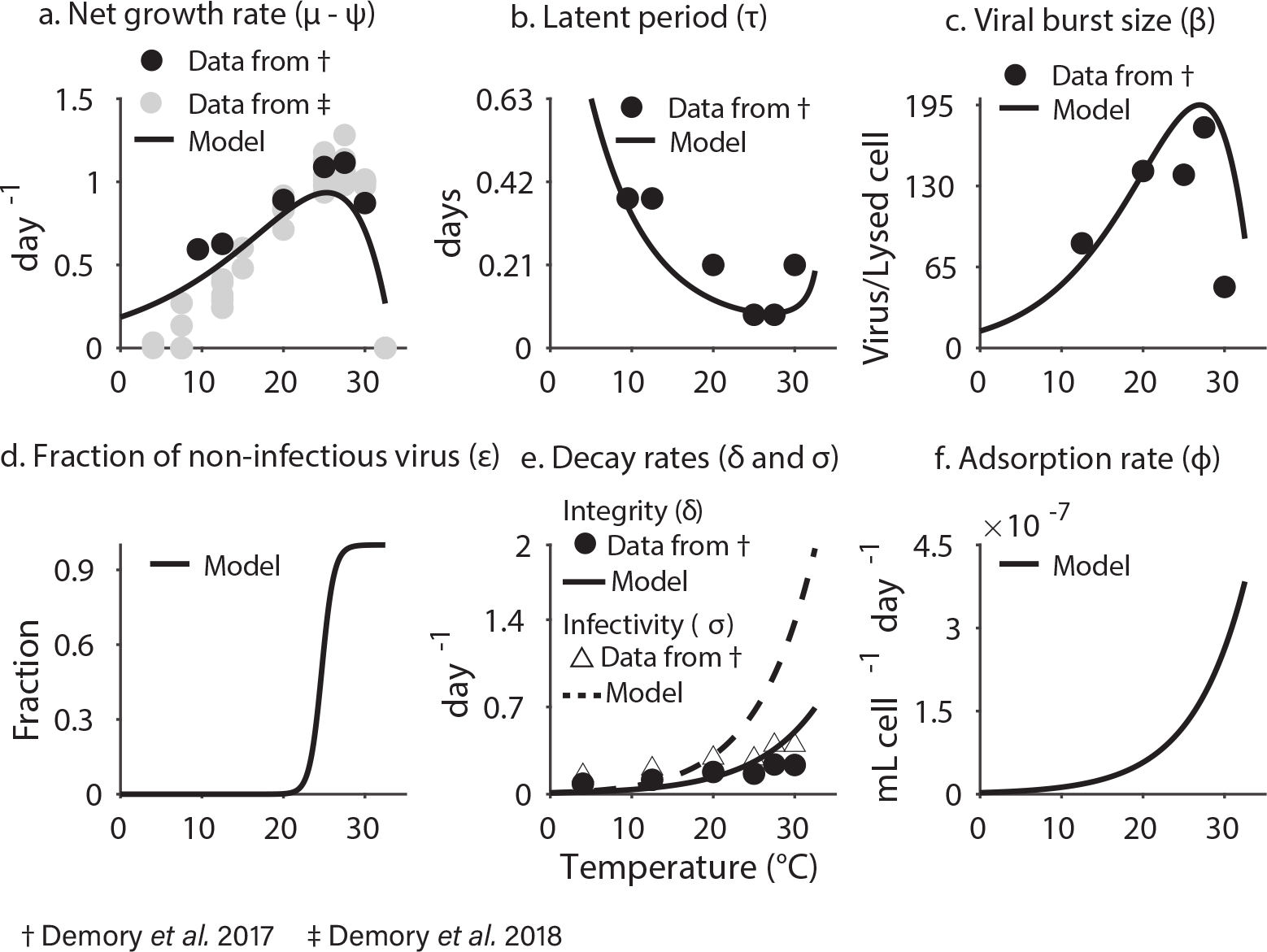
Parameters as functions of temperature. Black lines represent the parameter function used in the model. Black dots are the value of the parameter calculated experimentally in (Demory *et al*. 2017). Grey dots (top-left panel) are the growth rate values calculated experimentally in Demory *et al*. (2018).

### Viral fitness is maximal at a sub-optimal host growth temperature

We looked at the impact of temperature on the virus ability to invade an environment at the disease-free equilibrium (DFE). To do so, we computed the basic reproductive number 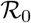 (see methods). Three different ecological states can be described (Fig. 4a): (1) when 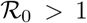, viruses can invade a susceptible host population. Depending on the temperature and the parameters set, the coexistence can be stable or unstable, and occurs only during a transition period leading to other states. We call this state the endemic state. (2) When 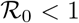, if the host can grow, the system reaches the DFE where viruses disappear from the medium. We call this phase the refuge state because the host is free from an epidemic. (3) If the host cannot grow, the system reaches the habitat loss state where neither the host nor virus can survive. We obtained the following condition for endemic coexistence (see methods):

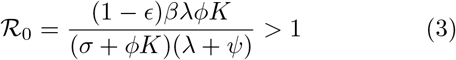

**FIG. 4:**
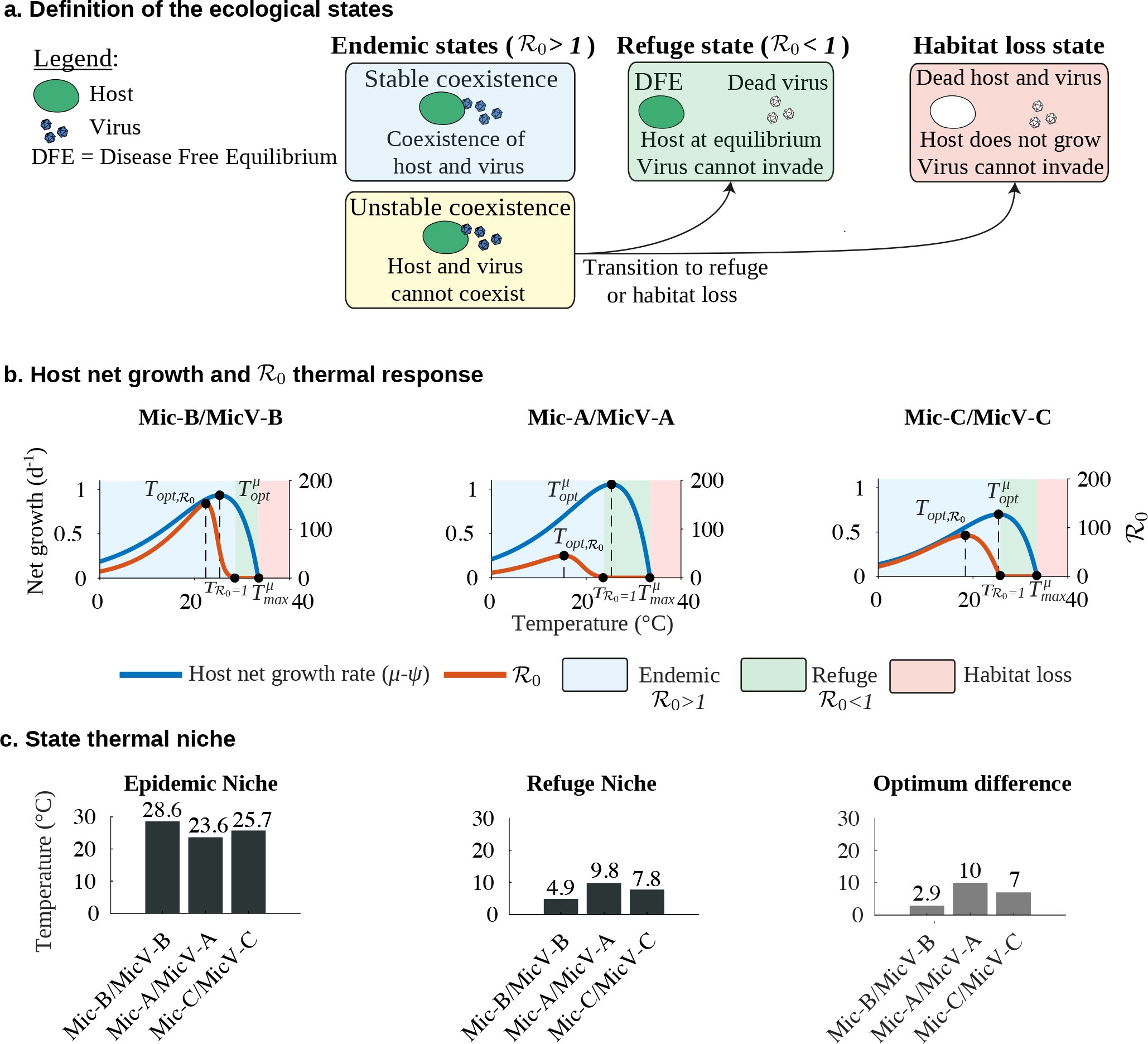
Ecological states as function of temperature. (a) Definition of the ecological states. (b) Host net growth (blue) vs. 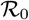 (orange) responses to temperature for the three host-virus systems. (c) Ecological thermal niches: endemic niche 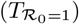, refuge niche 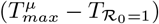 and optimum difference 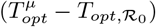.

This condition is the number of newly produced infectious cells given a single infected cell in the population (Li *et al*. 2019, Weitz *et al*. 2019). Invasion is possible when an infected cell leads to more than one infected cell on average (Weitz *et al*. 2019). The balance between viral production and removal followed the same thermal response for the three virus-host pairs (Fig. 4b). Endemic thermal niche (define as temperatures bellow 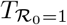) shifted at lower temperatures compared to the host thermal niches (define as temperatures between 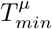 and 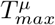) for the three pairs. This temperature shift was 6.6 ± 3.6°C on average. The endemic thermal niche (Fig. 4c) was 26.0± 2.5°C on average, whereas the refuge thermal niche was 7.5± 2.5°C on average. Pairs with a lower difference between optimal temperature of growth 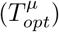 and optimal 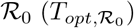 had a restricted thermal refuge niche but an extended endemic niche (Supplementary Fig. 5). Cardinal parameters for the three virus-host pairs are listed in Supplementary Tab. 4.

### Climate change impacts on ecological states and long-term host-virus coexistence

To generalize our previous analysis to global monthly averaged latitudinal SST projections (Fig. 5, see methods), we used our model for each virus-host pair for present (before 2020) to future periods (until 2100) based on dynamical monthly averaged SST from IPCC model GFCM2.1 ran with the SRES A2 scenario (GFDL Data Portal: http://nomads.gfdl.noaa.gov/). We then defined the ecological states based on the host-virus thermal niches (see methods). For the present period (Fig. 5a), the three virus-hosts pairs had relatively similar initial states based on their ecological thermal niche (Fig. 4). Due to the high 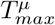 of the three hosts (Supplementary Tab. 2), we observed almost no habitat loss initially. Most of the ocean was in the endemic state with 94, 74 and 81% of its surface for Mic-B/MicV-B, Mic-A/MicV-A and Mic-C/MicV-C respectively. The refuge state was restricted to the tropics (30°S to 30°N). For future SST projections, we observed a notable transformation of the ecological state distribution (Fig. 5 b,c and d). The endemic area decreased for the three pairs with a loss of 13%, 7% and 9%, respectively, compared to the present period. The future refuge state of the three pairs increased by 12%, 6%, and 8% when compared to current conditions. Habitat loss only increased by 1% and was largely confined to the Red Sea (see Zoom-in in Fig. 5a and b). Mic-A/MicV-A and Mic-C/MicV-C pairs were more resilient than Mic-B/MicV-B, with a loss rate less than 10% of their endemic area (Fig. 5c) and a decrease rate lower than 0.01 percent per month. For all three virus-host pairs, the loss of the endemic zone occurred in the tropical and sub-tropical regions, where viruses live close to their maximal thermal limit 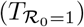.

**FIG. 5:**
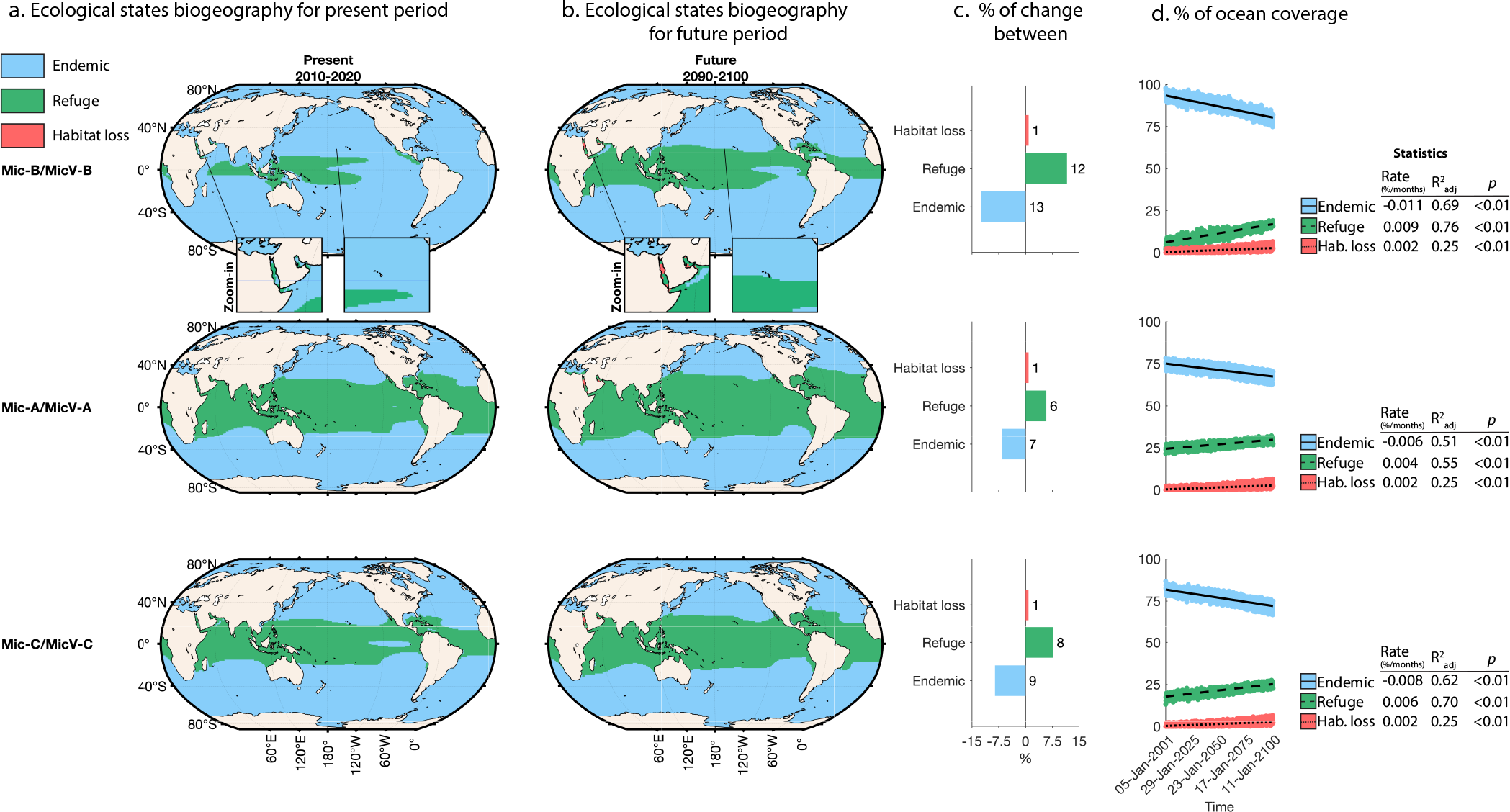
Ecological state biogeography between present (a) and future (b) periods given SST projection at global scale for the pairs Mic-B/MicV-B (first row), Mic-A/MicV-A (second row) and Mic-C/MicV-C (third row). The states are: endemic (host and virus coexistence – blue), refuge (host only – green) and habitat loss (host and virus cannot grow – red). (c) Coverage differences between present and future conditions in percentage of the ocean surface. (d) Percentage of ocean coverage dynamics from 2001 to 2100. Colored dots are model results for each month and black lines are linear regressions. The rates are calculated from a linear regression between the percentage of coverage and the time (each month).

## Discussion

We investigated the impact of temperature on the viral infection of *Micromonas* using data integration with a non-linear population model. We found that temperature-driven host and viral life-history traits explained population dynamics in the experiment for a broad thermal range. In doing so we found that incorporating non-infectious virus particles was critical to explain the observed population dynamics. Collectively, these findings suggest that the temperature-driven model captured essential processes that govern *Micromonas*-virus dynamics under different thermal conditions.

Our model-data integration shows that dynamics are driven by a temperature-dependent trade-off between virus production and loss of infectivity. This tradeoff suggests that increasing lysis and burst size generates higher amounts of non-infectious viruses. We are unaware of similar evidence in marine systems, however such a trade-off has been shown in phage infecting *Escherichia coli* (De Paepe & Taddei 2006). In this study, phage mortality was found to be proportional to the phage multiplication rate linked to viral capsid thickness and diameter. In other words, viruses produced in larger quantities may be less stable due to a thinner capsid. More generally, trade-offs may be important in understanding the link between viral particle size and other life-history traits such as burst size, latent period, and decay (Edwards *et al*. 2020). Our study also highlights the critical role between viral life-history trait trade-offs and host-virus dynamics. This trade-off between viral production and degradation was found in the three virus-phytoplankton systems used in this study. Additional research is needed to link temperature and virus size with the production-degradation trade-off across a broad set of host-virus pairs.

As described in the Results, the temperature-dependent model included a key life history trait: the fraction of non-infectious viruses produced during a burst. We find that including this term is critical to understand the observed dynamics. Specifically, the non-infectious viruses fraction dominated the viral population beyond 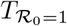 (Supplementary Fig. 6), supporting the importance of investigating infectivity both in experiments and in the field. Previous studies highlighted the loss of viral infectivity with temperature (Brown *et al*. 2006, Demory *et al*. 2017) and the small number of infected hosts (Baran *et al*. 2018) in warm environments. However, other mechanisms such as host resistance (Bellec *et al*. 2014, Clerissi *et al*. 2012, Thomas *et al*. 2011) can be responsible for losses of infectivity at higher temperatures and may explain the increased production of non-infectious viruses that we observed. Results from virus-prasinophyte systems have not shown significant relationships between host resistance and viral life-history traits (Ruiz *et al*. 2017) nor the impact of high temperature on resistance (Heath & Collins 2016). Given that other studies have measured resistance for up to 6 days in a similar biological system (e.g. *Ostreococcus*-Prasinovirus, (Yau *et al*. 2018)), the fraction of resistant cells in our study may be small given the 5-day duration of the experiment (Demory *et al*. 2017). Altogether, our findings suggest that loss of infectivity at high temperatures may alter the structure and function of the new viruses even when viruses are still in the host cell. More studies are needed to investigate the interplay between phenotypic and genotypic resistance from the host and the loss of infectivity of viral particles.

In addition to viral life-history traits, temperature also drives viral fitness and viral ecological niche. It is challenging to quantify viral fitness, especially in aquatic parasite-host systems. Using the epidemiological concept of 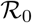 allowed us to quantify the role of temperature on viral invasion fitness. The comparison between 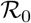 and host net growth thermal niches showed that viral invasion fitness does not have the same temperature response as host growth. We speculate that variations in thermal epidemic niches among viruses may also provide advantages at different temperatures (Supplementary Fig. 7). Viral fitness thermal curves may also reflect adaptation to the thermal host response. A virus that is well adapted to its host may have an optimal fitness temperature closely aligned to its host optimal temperature (Supplementary Fig. 5). Virus-host co-evolution may force phytoplankton to adapt to warmer temperatures with adaptation pressures from viruses at lower temperatures, and from deleterious impacts of high temperatures (Uiterwaal *et al*. 2020) (an issue we return to later in the Discussion).

A central objective of this study was to examine how variation in temperature can modulate virus-host coexistence. Host and virus long-term dynamics have been observed in the environment (Arsenieff *et al*. 2020, Baudoux *et al*. 2015, Brum *et al*. 2016, Vardi *et al*. 2012) and in laboratory-based experiments (Frickel *et al*. 2016, Marston *et al*. 2012, Shapiro *et al*. 2010, Thyrhaug *et al*. 2003). Coexistence between host and virus has often been explained by host resistance or phenological successions in marine communities (Avrani *et al*. 2011, Thyrhaug *et al*. 2003, Yau *et al*. 2020). Our model also allows for long-term coexistence at the population level. Temperature variations impact long-term coexistence at temperatures close to the maximal temperature of the endemic zone 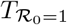 (Supplementary Fig. 10). This phenomenon is due to the nonlinear thermal response of the *Micromonas-virus* system. Temperature variations around 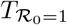 allowed the succession of favorable 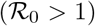 and unfavorable 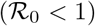 endemic conditions. On the one hand, these variations drove the loss of the virus for the average constant temperatures. This process was increased by a combination of long unfavorable periods beyond 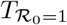 and high temperature variations. On the other hand, when the average temperatures were slightly higher than 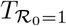 (unfavorable condition), variations allowed for coexistence of host and viruses. During the favorable periods, the virus reproduced efficiently allowing the maintenance of virus particles in the environment during unfavorable periods. The impact of fluctuating temperatures on the long-term dynamics depends on 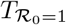 that is strain specific. We conclude that temperature fluctuations will have different impacts depending on the virus-host pair, leading to complex interactions at the community level, with different outcomes for phytoplankton competition and diversity (Chesson & Kuang 2008), and potentially influencing seasonal viral successions (Hevroni *et al*. 2020).

Finally, we extended our results on invasion and coexistence to large spatiotemporal scales. Using SST projections, we showed that ocean warming may have a significant impact on host-virus dynamics at a global scale (Fig. 5). Warming could reduce endemic zones – mainly in the tropics – where temperatures are already high. This pattern is similar for the three systems tested in our study but differs in terms of magnitude. The Mic-B/MicV-B pair is less resilient to warming compared to the two other pairs. This is due to its higher *T_opt,R0_*, which means that its distribution is limited to certain endemic zones in the tropics. In contrast, the two other pairs exhibit lower *T_opt,R0_* and have refuge states throughout the tropics. At the same time, hosts used in this study have a high 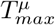 (Supplementary Tab. 2), which could mitigate habitat losses that would be expected due to increasing temperatures. Other subtropical and temperate hosts with lower 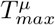 may be less resilient. Thus, our results suggest that the responses of host-virus pairs to climate change may vary considerably depending on the species involved. By considering viruses from this study, warming of the ocean may lead to more refuge environments in the tropics. Our model does not take into account host and virus evolution, which are an essential feature of microbial dynamics. The complexity of host-virus interactions are compounded by the limited data available, and necessitated the development of a simple model. However, our study suggests that evolution or adaptation processes will be necessary for virus and their host to coexist in warmer environments. These processes may drive dynamic shifts of their temperature-driven traits. Understanding eco-evolutionary dynamics of viruses and microbes is critical as the impacts of ocean warming increase in prevalence and severity (Gattuso *et al*. 2015, Hoegh-Guldberg *et al*. 2018, Reichert *et al*. 2002, Woodward *et al*. 2014).

## Material and Methods

### Laboratory population dynamics

To parameterize our model, we used population dynamics in laboratory from Demory *et al*. (2017). Three biological virus-host pairs were used in this study: Mic-A/MicV-A (RCC451/RCC4253), Mic-B/MicV-B (RCC829/RCC4265) and Mic-C/MicV-C (RCC834/RCC4229). Briefly, host and virus were obtained from the Roscoff Culture Collection. Infection experiments were performed at five or six temperature (9.5, 12.5, 20, 25, 27.5 and 30°C). Cultures were grown in batch conditions in a K-Si medium under the same light cycle (12:12 light/dark with 100 *μ*mol photons m^−2^ s^−1^) after a phase of acclimation of at least one month. Host and virus counts were performed by flow cytometry, every 4 hours for 120 hours. For more details on the experimental setup, see Demory *et al*. (2017).

### Life-history trait temperature-driven functions

#### Host growth and degradation

In line with the work of Grimaud (2016), the Hinshelwood model (Hinshelwood 1946) was used to represent the impact of temperature on gross phytoplankton growth (*μ*) and non-lysis mortality (*ψ*) rates as follows:

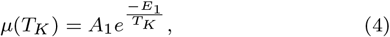

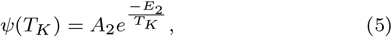

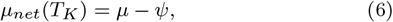

Here, *T_K_* is the temperature in Kelvin and parameters *A*_1_, *A*_2_, *E*_1_ and *E*_2_ are constants. Subtracting the two exponential (eq. 6), the net growth rate presents the typical phytoplankton response to temperature (Ras *et al*. 2013), with an asymmetric shape (Supplementary Fig. 8). This function is accurately represented by four empirical parameters when linked to the BR model (Bernard & Rémond 2012): the optimal growth temperature 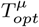 at which growth is maximal (*μ* = *μ_opt_*), and the minimal and maximal temperatures of growth (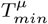 and 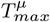 respectively). We express cardinal parameters form the Hinshelwood model according to Grimaud (2016):

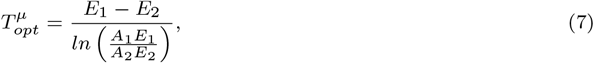

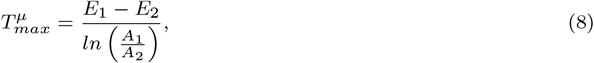

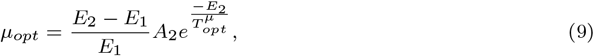

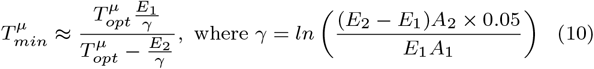

Finally, we hypothesized that the carrying capacity *K* did not depend on temperature and set *K* = 1.10^9^ cell.L^−1^ based on the work in Demory *et al*. (2018). This value is consistent with the experiments conducted under non-limiting nutrients, and in which measurements were performed during the exponential growth phase.

#### Thermal threshold between viral production and loss of infectivity

We defined the lysis rate as λ = 1/*τ*, where *τ* is the latent period, the time between the adsorption of the viral genome into the cell and the lysis of the host cell (Weitz 2016). To discriminate the mechanisms of the production and degradation of viruses, we described the temperature-driven lysis rate with a Hinshelwood model (Hinshelwood 1946):

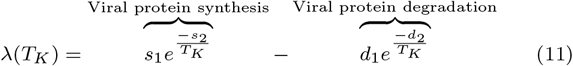

where *T_K_* is the temperature in Kelvin and parameters *s*_1_, *s*_2_, *d*_1_ and *d*_2_ are constants. The first exponential describes viral protein synthesis, whereas the second exponential describes viral protein degradation.

Burst size *β* is the number of viruses released per lysed host cell (Weitz 2016). The temperature may mainly act on the eclipse period (the time from adsorption to detection of an assembled virus in the cell), explaining the temperature-driven dynamics of burst size. Therefore, the burst size turned out to be efficiently represented by the same temperature function as the lysis rate:

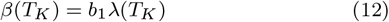

where *b*_1_ is a scaling coefficient.

The viral loss of infectivity is likely to be due to protein degradation, viral membrane fluidity or structural modification when the temperature increases. We then represented the loss of infectivity with two parameters: the production of non-infectious viruses by lysed cell and loss of infectivity outside the host cell. We described this process as a logistic function with a rate proportional to the viral protein degradation (eq. 11):

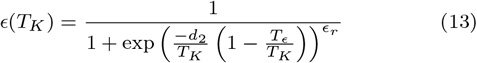

with *d*_2_/*T_K_* is the rate of protein degradation from eq. 11, *T_ϵ_* is the temperature where 50% of the produced viruses are non-infectious and *ϵ_r_* is a scaling constant.

The second process of virus loss of infectivity is the degradation of viruses in the medium. In line with De Paepe & Taddei (2006), Demory *et al*. (2017) we represented this degradation as an Arrhenius function:

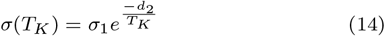

where 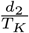 is the rate of protein degradation from eq. 11 and *σ*_1_ is a scaling coefficient.

Similarly to the loss of infectivity, viruses in the medium decay at an exponential rate with temperature (De Paepe & Taddei 2006, Demory *et al*. 2017):

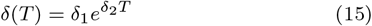

#### Viral adsorption

Adsorption rate *ϕ* is a process that depends on particle movements (virus probability to encounter a host cell) and on the specific receptors on the host and virus membrane (Weitz 2016). Murray and Jackson have mechanically defined this parameter based on physical considerations (Murray & Jackson 1992). The maximum adsorption rate is given by 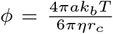, where *a* is the host cell radius, *k_b_* the Boltzmann constant, *η* the viscosity at the absolute temperature *T* and *r_c_* the viral particle radius (Murray & Jackson 1992, Talmy *et al*. 2019). We do not have evidence that host and virus radius are affected by temperature in this range (Supplementary Fig. 9), and therefore *η* is the temperature-driven parameter. The shear viscosity of water *η* decreases with temperature at an exponential rate (Reynolds 1886). Therefore, *ϕ* turns out to be increasing with temperature. We then represented *ϕ* by the following function:

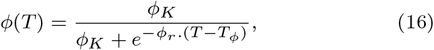

where *T_ϕ_* is the half temperature saturation, *ϕ_r_* is the exponential rate, and *ϕ_K_* is the saturating adsorption value.

### Model calibration

The calibration procedure proceeds in three steps. First, basal model parameters were identified for each experiment at a set temperature (*T_i_*) independently, based on experimental measurements of both hosts (*H_data_*) and free viruses (*V_data_*). For any given parameter set (*θ*), model predictions can be computed at each measurement time (*t_j_*) for both host (*H_model_*) and virus (*V_model_*). The calibration procedure consisted in minimizing the following error between predictions and measurements:

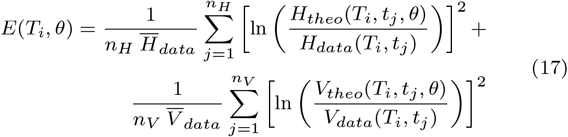

Where *n_H_* and *n_V_* are the number of measurements for the host and virus data respectively, and 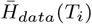 and 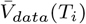 are the mean of the host and virus data respectively, determined for each temperature *T_i_*. We used a MATLAB Differential Evolution algorithm Yarpiz (2020) to identify the set of parameters *θ*\*(*T_i_*) that minimizes the error criteria *E*.

We then built temperature-driven functions for the ten basal model parameters. The second stage consisted in estimating, for each basal model parameter, the hyper-parameters set (*η_k_*) of the temperature-driven function *f_k_*(*T, η_k_*), based on the previous parameter estimations *θ*\*(*T_i_*). As parameters *K* and *ω* were assumed to be temperature-independent, we set constant functions for these two parameters. We computed the global error aggregating all experimental data for the different temperature as follows:

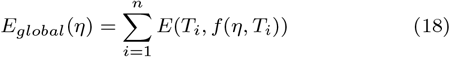

where *f* is the vector of the 10 temperature-driven functions and *η* the vector of the 19 hyper-parameters. The same optimization algorithm was run to determine the optimal hyper-parameter set *η*\*, minimizing this global error. The resulting optimal hyperparameters are given in Supplementary Tab. 3.

### Model fit representation

We represented the model fits with a thermal variation of ±1°C degree to account for the variability of the thermal laboratory system. Using a Monte-Carlo procedure, for each temperature *T_i_* we ran the model for 10,000 temperature values between *T_i_*-1 and *T_i_*+1. We then calculated the median simulation fit and its variation range.

### Quantifying model accuracy

To quantify the model accuracy to fit experimental data, we computed the Akaike information criterion (AIC) and the Bayesian information criterion (BIC) for the temperature-driven model and the basal model according to the following equations Burnham & Anderson (2004):

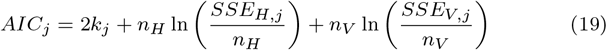

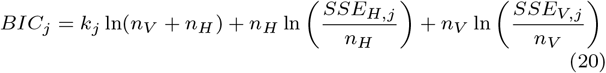

with

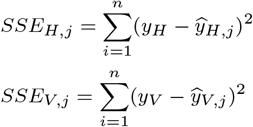

Where *k* is the number of parameters to be estimated (10 for the basal model and 19 for the temperature-driven model), *j* is the model, *i* is the data point, *n* the maximum number of data, *y_H_* and *y_V_* are the host and virus data and 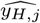 and 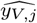 are the host and virus estimations for model *j*.

### Basic reproductive number

To explore the impact of temperature on the viral infection dynamics, we computed the basic reproductive number 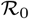, defined as the number of secondary infections resulting from the introduction of an infectious individual into a entirely susceptible population (Anderson & May 1982). We calculated this criterion according to Weitz *et al*. (2019) by considering *V_i_* the epidemiological birth state from equation 2. The terms for a new infections are: (1 – *ϵ*)*β*λ. The loss terms are: *ψI* and *ϕSV_i_* + *σV_i_* + *ω*(*V*)*V_i_*. By replacing *I* by 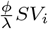 and considering the system at DFE (K,0,0,0) with only a susceptible host population we obtained:

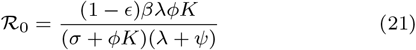

### Global temperature data and analysis

We used monthly averaged SST projections from the IPCC model GFCM2.1 ran with the SRES A2 scenario (GFDL Data Portal: http://nomads.gfdl.noaa.gov/) from 2001 to 2100. We defined two periods: the present period from 2010 to 2020 and the future period from 2090 to 2100. To calculate the ecological state (endemic, refuge, or habitat loss) for each location of the ocean, we compare the thermal niche of the three virus-host pairs to the averaged ten years SST (*T_i_*) for the period *i* (present or future) as follows:

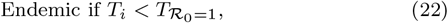

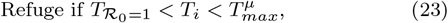

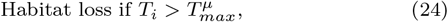

where 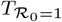 is the maximal temperature of the endemic niche and 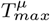 is the maximal temperature of the host net growth. We then calculated the differences between present and future states as the differences in percentage of ocean coverage between present and future periods.

## Authorship statement

D.D., S.T., and O.B. designed the study. D.D., O.B. and J.S.W. contributed to model design and analysis. D.D. provided the display items. A-C.B., N.S., A.S. and S.R. contributed to the technical and biological design of the project. D.D., J.S.W and O.B. wrote the manuscript with contributions from all authors. All authors approved the final manuscript.

## Data accessibility statementme

The data and code supporting the results are accessible at: https://zenodo.org/badge/latestdoi/288514967.

## Acknowledgments

We are particularly grateful to Cody Clements, Yves Desdevises, Guanlin Li, Rozenn Pineau and Andreea Magalie for providing comments about the manuscript and the model. We also thank members of the Ecology of Marine Plankton team (Biological Station of Roscoff), the Roscoff Culture Collection, the INRIA BIOCORE team and the Weitz group at Georgia Institute of Technology for our discussions and their insights on our study.

## References

Abedon, S.T. (2008). Bacteriophage ecology: population growth, evolution, and impact of bacterial viruses. vol. 15. Cambridge University Press.

Anderson, R.M. & May, R.M. (1982). Population Biology of Infectious Diseases: Report of the Dahlem Workshop on Population Biology of Infectious Disease Agents, Berlin 1982, March 14-19. Springer.

Arsenieff, L., Le Gall, F., Rigaut-Jalabert, F., Mahé, F., Sarno, D., Gouhier, L., Baudoux, A.C. & Simon, N. (2020). Diversity and dynamics of relevant nanoplanktonic diatoms in the western English Channel. The ISME Journal.

Avrani, S., Wurtzel, O., Sharon, I., Sorek, R. & Lindell, D. (2011). Genomic island variability facilitates *Prochlorococcus*-virus coexistence. Nature, 474, 604–608.

Baran, N., Goldin, S., Maidanik, I. & Lindell, D. (2018). Quantification of diverse virus populations in the environment using the Polony method. Nature Microbiology, 3, 62.

Baudoux, A.C. & Brussaard, C.P. (2005). Characterization of different viruses infecting the marine harmful algal bloom species *Phaeocystis globosa*. Virology, 341, 80–90.

Baudoux, A.C., Lebredonchel, H., Dehmer, H., Latimier, M., Edern, R., Rigaut-Jalabert, F., Ge, P., Guillou, L., Foulon, E., Bozec, Y. et al. (2015). Interplay between the genetic clades of m icromonas and their viruses in the western english channel. Environmental Microbiology Reports, 7, 765–773.

Baudoux, A.C., Veldhuis, M.J., Witte, H.J. & Brussaard, C.P. (2007). Viruses as mortality agents of picophytoplankton in the deep chlorophyll maximum layer during ironages iii. Limnology and Oceanography, 52, 2519–2529.

Béchette, A., Stojsavljevic, T., Tessmer, M., Berges, J.A., Pinter, G.A. & Young, E.B. (2013). Mathematical modeling of bacteria-virus interactions in lake Michigan incorporating phosphorus content. Journal of Great Lakes Research, 39, 646–654.

Bellec, L., Clerissi, C., Edern, R., Foulon, E., Simon, N., Grimsley, N. & Desdevises, Y. (2014). Cophylogenetic interactions between marine viruses and eukaryotic picophytoplankton. BMC Evolutionary Biology, 14, 59.

Bernard, O. & Réemond, B. (2012). Validation of a simple model accounting for light and temperature effect on microalgal growth. Bioresource Technology, 123, 520–527.

Bohannan, B.J. & Lenski, R.E. (1997a). Effect of resource enrichment on a chemostat community of bacteria and bacteriophage. Ecology, 78, 2303–2315.

Bohannan, B.J. & Lenski, R.E. (1997b). Effect of resource enrichment on a chemostat community of bacteria and bacteriophage. Ecology, 78, 2303–2315.

Bratbak, G., Jacobsen, A., Heldal, M., Nagasaki, K. & Thingstad, F. (1998). Virus production in *Phaeocystis pouchetii* and its relation to host cell growth and nutrition. Aquatic Microbial Ecology, 16, 1–9.

Breitbart, M., Bonnain, C., Malki, K. & Sawaya, N.A. (2018). Phage puppet masters of the marine microbial realm. Nature Microbiology, 3, 754–766.

Brown, C.M., Lawrence, J.E. & Campbell, D.A. (2006). Are phytoplankton population density maxima predictable through analysis of host and viral genomic DNA content? Journal of the Marine Biological Association of the United Kingdom, 86, 491–498.

Brum, J.R., Hurwitz, B.L., Schofield, O., Ducklow, H.W. & Sullivan, M.B. (2016). Seasonal time bombs: dominant temperate viruses affect southern ocean microbial dynamics. The ISME Journal, 10, 437–449.

Burnham, K.P. & Anderson, D.R. (2004). Multimodel inference: understanding aic and bic in model selection. Sociological methods & research, 33, 261–304.

Chesson, P. & Kuang, J.J. (2008). The interaction between predation and competition. Nature, 456, 235–238.

Clerissi, C., Desdevises, Y. & Grimsley, N. (2012). Prasinoviruses of the marine green alga *Ostreococcus tauri* are mainly species specific. Journal of Virology, 86, 4611–4619.

Cordova, A., Deserno, M., Gelbart, W.M. & Ben-Shaul, A. (2003). Osmotic shock and the strength of viral capsids. Biophysical Journal, 85, 70–74.

De Paepe, M. & Taddei, F. (2006). Viruses’ life history: towards a mechanistic basis of a trade-off between survival and reproduction among phages. PLOS Biology, 4, e193.

Demory, D., Arsenieff, L., Simon, N., Six, C., Rigaut-Jalabert, F., Marie, D., Ge, P., Bigeard, E., Jacquet, S., Sciandra, A. et al. (2017). Temperature is a key factor in *Micromonas* virus interactions. The ISME Journal, 11, 601.

Demory, D., Baudoux, A.C., Monier, A., Simon, N., Six, C., Ge, P., Rigaut-Jalabert, F., Marie, D., Sciandra, A., Bernard, O. & Rabouille, S. (2018). Picoeukaryotes of the *Micromonas* genus: sentinels of a warming ocean. The ISME Journal.

Edwards, K.F., Steward, G.F. & Schvarcz, C.A. (2020). Making sense of virus size and the tradeoffs shaping viral fitness. bioRxiv.

Frickel, J., Sieber, M. & Becks, L. (2016). Eco-evolutionary dynamics in a coevolving host-virus system. Ecology letters, 19, 450–459.

Fuhrman, J.A. (1999). Marine viruses and their biogeochemical and ecological effects. Nature, 399, 541–548.

Gattuso, J.P., Magnan, A., Billé, R., Cheung, W.W., Howes, E.L., Joos, F., Allemand, D., Bopp, L., Cooley, S.R., Eakin, C.M. et al. (2015). Contrasting futures for ocean and society from different anthropogenic CO2 emissions scenarios. Science, 349, aac4722.

Gerla, D.J., Gsell, A.S., Kooi, B.W., Ibelings, B.W., Van Donk, E. & Mooij, W.M. (2013). Alternative states and population crashes in a resource-susceptible-infected model for planktonic parasites and hosts. Freshwater Biology, 58, 538–551.

Grimaud, G.M. (2016). Modelling the temperature effect on phytoplankton: from acclimation to adaptation. Ph.D. thesis, Université Nice Sophia Antipolis; France.

Heath, S.E. & Collins, S. (2016). Mode of resistance to viral lysis affects host growth across multiple environments in the marine picoeukaryote *Ostreococcus tauri*. Environmental Microbiology, 18, 4628–4639.

Hevroni, G., Flores-Uribe, J., Béjà, O. & Philosof, A. (2020). Rising through the ranks: Seasonal and diel patterns of marine viruses. bioRxiv.

Hinshelwood, C.N. (1946). Chemical kinetics of the bacterial cell. The Clarendon Press; London.

Hoegh-Guldberg, O., Jacob, D., Taylor, M., Bindi, M., Brown, S., Camilloni, I., Diedhiou, A., Djalante, R., Ebi, K., Engelbrecht, F. et al. (2018). Impacts of 1.5 °C global warming on natural and human systems. Intergovernmental Panel on Climate Change.

Jacquet, S., Miki, T., Noble, R., Peduzzi, P. & Wilhelm, S. (2010). Viruses in aquatic ecosystems: important advancements of the last 20 years and prospects for the future in the field of microbial oceanography and limnology. Advances in Oceanography and Limnology, 1, 97–141.

Jover, L.F., Effler, T.C., Buchan, A., Wilhelm, S.W. & Weitz, J.S. (2014). The elemental composition of virus particles: implications for marine biogeochemical cycles. Nature Reviews Microbiology, 12, 519.

Kendrick, B.J., DiTullio, G.R., Cyronak, T.J., Fulton, J.M., Van Mooy, B.A. & Bidle, K.D. (2014). Temperature-induced viral resistance in *Emiliania huxleyi* (prymnesiophyceae). PLOS ONE, 9, e112134.

Knowles, B., Silveira, C., Bailey, B., Barott, K., Cantu, V., Cobian-Guemes, A., Coutinho, F., Dinsdale, E., Felts, B., Furby, K. et al. (2016). Lytic to temperate switching of viral communities. Nature, 531, 466.

Li, G., Cortez, M.H., Dushoff, J. & Weitz, J.S. (2019). When to be temperate: On the fitness benefits of lysis vs. lysogeny. Virus Evolution.

Marston, M.F., Pierciey, F.J., Shepard, A., Gearin, G., Qi, J., Yandava, C., Schuster, S.C., Henn, M.R. & Martiny, J.B. (2012). Rapid diversification of coevolving marine *Synechococcus* and a virus. Proceedings of the National Academy of Sciences, 109, 4544–4549.

Martin-Platero, A.M., Cleary, B., Kauffman, K., Preheim, S.P., McGillicuddy, D.J., Alm, E.J. & Polz, M.F. (2018). High resolution time series reveals cohesive but short-lived communities in coastal plankton. Nature communications, 9, 1–11.

Martínez, J.M., Boere, A., Gilg, I., van Lent, J.W., Witte, H.J., van Bleijswijk, J.D. & Brussaard, C.P. (2015). New lipid envelope-containing dsDNA virus isolates infecting *Micromonas pusilla* reveal a separate phylogenetic group. Aquatic Microbial Ecology, 74, 17–28.

Middelboe, M. (2000). Bacterial growth rate and marine virus-host dynamics. Microbial Ecology, 40, 114–124.

Middelboe, M., Hagström, A., Blackburn, N., Sinn, B., Fischer, U., Borch, N., Pinhassi, J., Simu, K. & Lorenz, M. (2001). Effects of bacteriophages on the population dynamics of four strains of pelagic marine bacteria. Microbial Ecology, 42, 395406.

Mojica, K.D. & Brussaard, C.P. (2014). Factors affecting virus dynamics and microbial host-virus interactions in marine environments. FEMS microbiology ecology, 89, 495–515.

Mojica, K.D., Huisman, J., Wilhelm, S.W. & Brussaard, C.P. (2016). Latitudinal variation in virus-induced mortality of phytoplankton across the North Atlantic ocean. The ISME Journal, 10, 500.

Murray, A.G. & Jackson, G.A. (1992). Viral dynamics: a model of the effects of size, shape, motion and abundance of single-celled planktonic organisms and other particles. Marine Ecology Progress Series, pp. 103–116.

Nagasaki, K. & Yamaguchi, M. (1998). Intra-species host specificity of HaV *(Heterosigma akashiwo* virus) clones. Aquatic Microbial Ecology, 14, 109–112.

Perelson, A.S. (2002). Modelling viral and immune system dynamics. Nature Reviews Immunology, 2, 28.

Ras, M., Steyer, J.P. & Bernard, O. (2013). Temperature effect on microalgae: a crucial factor for outdoor production. Reviews in Environmental Science and Bio/Technology, 12, 153–164.

Reichert, B.K., Schnur, R. & Bengtsson, L. (2002). Global ocean warming tied to anthropogenic forcing. Geophysical Research Letters, 29, 20–1.

Reynolds, O. (1886). On the theory of lubrication and its application to Mr. Beauchamp tower’s experiments, including an experimental determination of the viscosity of olive oil. Philosophical Transactions of the Royal Society of London, 177, 157–234.

Rhodes, C. & Martin, A. (2010). The influence of viral infection on a plankton ecosystem undergoing nutrient enrichment. Journal of Theoretical Biology, 265, 225–237.

Rhodes, C., Truscott, J. & Martin, A. (2008). Viral infection as a regulator of oceanic phytoplankton populations. Journal of Marine Systems, 74, 216–226.

Ruiz, E., Baudoux, A.C., Simon, N., Sandaa, R.A., Thingstad, T.F. & Pagarete, A. (2017). *Micromonas* versus virus: New experimental insights challenge viral impact. Environmental Microbiology, 19, 2068–2076.

Samanta, S., Dhar, R., Pal, J. & Chattopadhyay, J. (2013). Effect of enrichment on plankton dynamics where phytoplankton can be infected from free viruses. Nonlinear Studies, 20.

Shapiro, O.H., Kushmaro, A. & Brenner, A. (2010). Bacteriophage predation regulates microbial abundance and diversity in a full-scale bioreactor treating industrial wastewater. The ISME Journal, 4, 327–336.

Suttle, C.A. (2005). Viruses in the sea. Nature, 437, 356.

Suttle, C.A. (2007). Marine viruses-major players in the global ecosystem. Nature Reviews Microbiology, 5, 801.

Talmy, D., Beckett, S.J., Taniguchi, D.A., Zhang, A.B., Weitz, J.S. & Follows, M.J. (2019). Contrasting controls on microzooplankton grazing and viral infection of microbial prey. Frontiers in Marine Science, 6, 182.

Thingstad, T.F. (2000). Elements of a theory for the mechanisms controlling abundance, diversity, and biogeochemical role of lytic bacterial viruses in aquatic systems. Limnology and Oceanography, 45, 1320–1328.

Thomas, R., Grimsley, N., Escande, M.l., Subirana, L., Derelle, E. & Moreau, H. (2011). Acquisition and maintenance of resistance to viruses in eukaryotic phytoplankton populations. Environmental Microbiology, 13, 1412–1420.

Thyrhaug, R., Larsen, A., Thingstad, T.F. & Bratbak, G. (2003). Stable coexistence in marine algal host-virus systems. Marine Ecology Progress Series, 254, 27–35.

Tomaru, Y., Kimura, K. & Yamaguchi, H. (2014). Temperature alters algicidal activity of DNA and RNA viruses infecting chaetoceros tenuissimus. Aquatic Microbial Ecology, 73, 171–183.

Tomaru, Y., Tanabe, H., Yamanaka, S. & Nagasaki, K. (2005). Effects of temperature and light on stability of microalgal viruses, hav, hcv and hcrnav. Plankton Biology and Ecology, 52, 1–6.

Uiterwaal, S.F., Lagerstrom, I.T., Luhring, T.M., Salsbery, M.E. & DeLong, J.P. (2020). Trade-offs between morphology and thermal niches mediate adaptation in response to competing selective pressures. Ecology and Evolution, 10, 1368–1377.

Vardi, A., Haramaty, L., Van Mooy, B.A., Fredricks, H.F., Kimmance, S.A., Larsen, A. & Bidle, K.D. (2012). Host-virus dynamics and subcellular controls of cell fate in a natural coccolithophore population. Proceedings of the National Academy of Sciences, 109, 19327–19332.

Weinbauer, M.G., Brettar, I. & Höofle, M.G. (2003). Lysogeny and virus-induced mortality of bacterioplankton in surface, deep, and anoxic marine waters. Limnology and Oceanography, 48, 1457–1465.

Weinbauer, M.G. & Rassoulzadegan, F. (2004). Are viruses driving microbial diversification and diversity? Environmental Microbiology, 6, 1–11.

Weinbauer, M.G., Wilhelm, S.W., Suttle, C.A., Pledger, R.J. & Mitchell, D.L. (1999). Sunlight-induced DNA damage and resistance in natural viral communities. Aquatic Microbial Ecology, 17, 111–120.

Weitz, J.S. (2016). Quantitative viral ecology: dynamics of viruses and their microbial hosts. Princeton University Press.

Weitz, J.S., Li, G., Gulbudak, H., Cortez, M.H. & Whitaker, R.J. (2019). Viral invasion fitness across a continuum from lysis to latency. Virus evolution, 5, vez006.

Wigginton, K.R., Pecson, B.M., Sigstam, T., Bosshard, F. & Kohn, T. (2012). Virus inactivation mechanisms: impact of disinfectants on virus function and structural integrity. Environmental Science & Technology, 46, 12069–12078.

Wilhelm, S. & Suttle, C. (2000). Viruses as regulators of nutrient cycles in aquatic environments. Microbial biosystems: New frontiers, edited by: Bell, CR, Brylinsky, M., and Johnson-Green, P., Atlantic Canada Society of Microbial Ecology, pp. 551–556.

Wilhelm, S.W., Weinbauer, M.G., Suttle, C.A. & Jeffrey, W.H. (1998). The role of sunlight in the removal and repair of viruses in the sea. Limnology and Oceanography, 43, 586–592.

Williamson, S.J., Cary, S.C., Williamson, K.E., Helton, R.R., Bench, S.R., Winget, D. & Wommack, K.E. (2008). Lysogenic virus-host interactions predominate at deep-sea diffuse-flow hydrothermal vents. The ISME Journal, 2, 1112.

Woodward, A., Smith, K.R., Campbell-Lendrum, D., Chadee, D.D., Honda, Y., Liu, Q., Olwoch, J., Revich, B., Sauerborn, R., Chafe, Z. et al. (2014). Climate change and health: on the latest IPCC report. The Lancet, 383, 1185–1189.

Yarpiz (2020). Differential evolution (de). https://www.mathworks.com/matlabcentral/fileexchange/52897-differential-evolution-de. MATLAB Central File Exchange.

Yau, S., Caravello, G., Fonvieille, N., Desgranges, E., Moreau, H. & Grimsley, N. (2018). Rapidity of genomic adaptations to prasinovirus infection in a marine microalga. Viruses, 10, 441.

Yau, S., Krasovec, M., Benites, L.F., Rombauts, S., Groussin, M., Vancaester, E., Aury, J.M., Derelle, E., Desdevises, Y., Escande, M.L. et al. (2020). Virus-host coexistence in phytoplankton through the genomic lens. Science Advances, 6, eaay2587.

