## Supplementary Information for "A thermal trade-off between viral production and degradation drives phytoplankton-virus population dynamics"

### Supplementary Tables

**Tab. S 1.** Information about variables and parameters used in the model

| Variables | Signification | Units |
|---|---|---|
| $S$ | susceptible phytoplankton | cell ml$^{-1}$ |
| $I$ | infected phytoplankton | cell ml$^{-1}$ |
| $V_i$ | free infectious viruses | virus ml$^{-1}$ |
| $V_{ni}$ | free non-infectious viruses | virus ml$^{-1}$ |
| **Parameters** | **Signification** | **Units** |
| $\mu$ | host gross growth rate | day$^{-1}$ |
| $\psi$ | host loss rate | day$^{-1}$ |
| $K$ | carrying capacity | cell ml$^{-1}$ |
| $\phi$ | adsorption rate | ml cell$^{-1}$ d$^{-1}$ |
| $\lambda$ | lysis rate | day$^{-1}$ |
| $\varepsilon$ | percentage of non-infectious viruses produced | unitless |
| $\beta$ | viral burst size | virus cell$^{-1}$ |
| $\sigma$ | loss of infectivity | day$^{-1}$ |
| $\delta$ | viral decay | day$^{-1}$ |
| $\omega$ | viral aggregation rate | ml virus$^{-1}$ d$^{-1}$ |

**Tab. S 2.** Akaike Information Criterion (AIC) and Bayesian Information Criterion (BIC) calculated for each temperature for the temperature-driven and basal models.

| Temperature | AIC | | BIC | |
|---|---|---|---|---|
| | Temperature-driven model | Basal model | Temperature-driven model | Basal model |
| 9.5 | **1177.9** | 1251.2 | **1210.4** | 1268.3 |
| 12.5 | **1298.3** | 1308.1 | 1331.3 | **1325.5** |
| 20 | **1134.2** | 1219 | **1166.3** | 1235.9 |
| 25 | **1206.6** | 1228.3 | **1238.7** | 1245.2 |
| 27.5 | **1229.6** | 1305 | 1331.7 | **1321.9** |
| 30 | **1241.7** | 1292.5 | **1274.3** | 1309.7 |

**Tab. S 3.** Hyper-parameter values of the temperature-driven functions.

| Hyper-parameters | Mic-B/MicV-B | Mic-A/MicV-A | Mic-C/MicV-C | Units |
|---|---|---|---|---|
| $A1$ | $1\ 10^{10}$ | $1.1748\ 10^{10}$ | $9.3729\ 10^{9}$ | $\text{day}^{-1}$ |
| $E1$ | 6745.1 | 6757.8 | 6812.3 | °K |
| $A2$ | $1\ 10^{25}$ | $1.1964\ 10^{25}$ | $8.5335\ 10^{24}$ | $\text{day}^{-1}$ |
| $E2$ | 17335 | 17354 | 17375 | °K |
| $K$ | $1\ 10^{9}$ | $1\ 10^{9}$ | $1.1\ 10^{7}$ | $\text{cell ml}^{-1}$ |
| $\phi_K$ | $1.2\ 10^{-6}$ | $1.037\ 10^{-6}$ | $1.03\ 10^{-6}$ | $\text{ml cell}^{-1}\ \text{day}^{-1}$ |
| $T_\phi$ | 40 | 47.235 | 60.251 | °C |
| $\phi_r$ | 0.15 | 0.057292 | 0.14717 | $°\text{C}^{-1}$ |
| $s1$ | $1.0138\ 10^{22}$ | $1.0137\ 10^{22}$ | $1.0159\ 10^{22}$ | $\text{day}^{-1}$ |
| $d1$ | 13119 | 13120 | 13121 | °K |
| $s2$ | $1.6256\ 10^{22}$ | $1.6265\ 10^{22}$ | $1.6421\ 10^{22}$ | $\text{day}^{-1}$ |
| $d2$ | 13264 | 13261 | 13265 | °K |
| $b1$ | 17 | 17.182 | 118.66 | $\text{virus day cell}^{-1}$ |
| $T_\varepsilon$ | 24.71 | 18.97 | 27.98 | °C |
| $\varepsilon_r$ | 8.937 | 4.291 | 3.00023 | unitless |
| $\sigma_1$ | $1.3859\ 10^{19}$ | $1.1388\ 10^{19}$ | $5.181\ 10^{18}$ | $\text{day}^{-1}$ |
| $\delta_1$ | 0.01 | 0.010065 | 0.000039574 | $\text{day}^{-1}$ |
| $\delta_2$ | 0.13046 | 0.00013046 | 0.060278 | $°\text{C}^{-1}$ |
| $\omega$ | $1.9217\ 10^{-8}$ | $1\ 10^{-8}$ | $8.9811\ 10^{-10}$ | $\text{ml virus}^{-1}\ \text{day}^{-1}$ |

**Tab. S 4.** Cardinal parameters for host growth and $\mathscr{R}_0$ for the three host-virus systems

| Parameters | Mic-B/MicV-B | Mic-A/MicV-A | Mic-C/MicV-C | Units |
|---|---|---|---|---|
| $T^{\mu}_{opt}$ | 25.30 | 25.33 | 25.382 | °C |
| $T^{\mu}_{max}$ | 33.46 | 33.47 | 33.497 | °C |
| $T^{\mu}_{min}$ | -10.33 | -10.35 | -10.438 | °C |
| $\mu_{opt}$ | 0.93 | 1.05 | 0.70 | $\text{day}^{-1}$ |
| $T_{opt,\mathscr{R}_0}$ | 22.38 | 15.32 | 18.375 | °C |
| $T_{\mathscr{R}_0=1}$ | 28.59 | 23.63 | 25.741 | °C |
| $\mathscr{R}_{0,opt}$ | 152.94 | 45.47 | 84.59 | unitless |

### Supplementary Figures

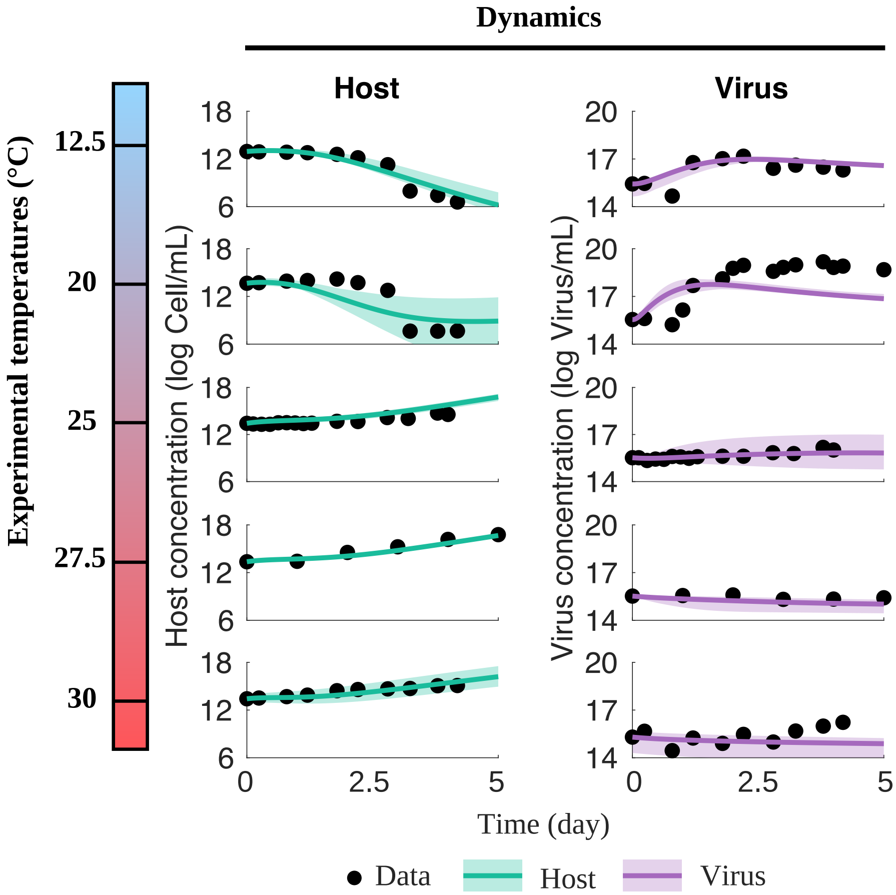


**Fig. S 1.** Model fits for the 5 temperatures tested experimentally in (1) for the pairs Mic-A/MicV-A (RCC451/RCC4263). The left column shows the phytoplankton dynamics with total host population $N$ model (green line) and experimental data (black circles). The right column shows the virus dynamics with total virus population $V$ model (purple line) experimental data (black circles). Shaded area represent the model fits variability when accounting for a $\pm$ 1°C uncertainty from experimental temperature.

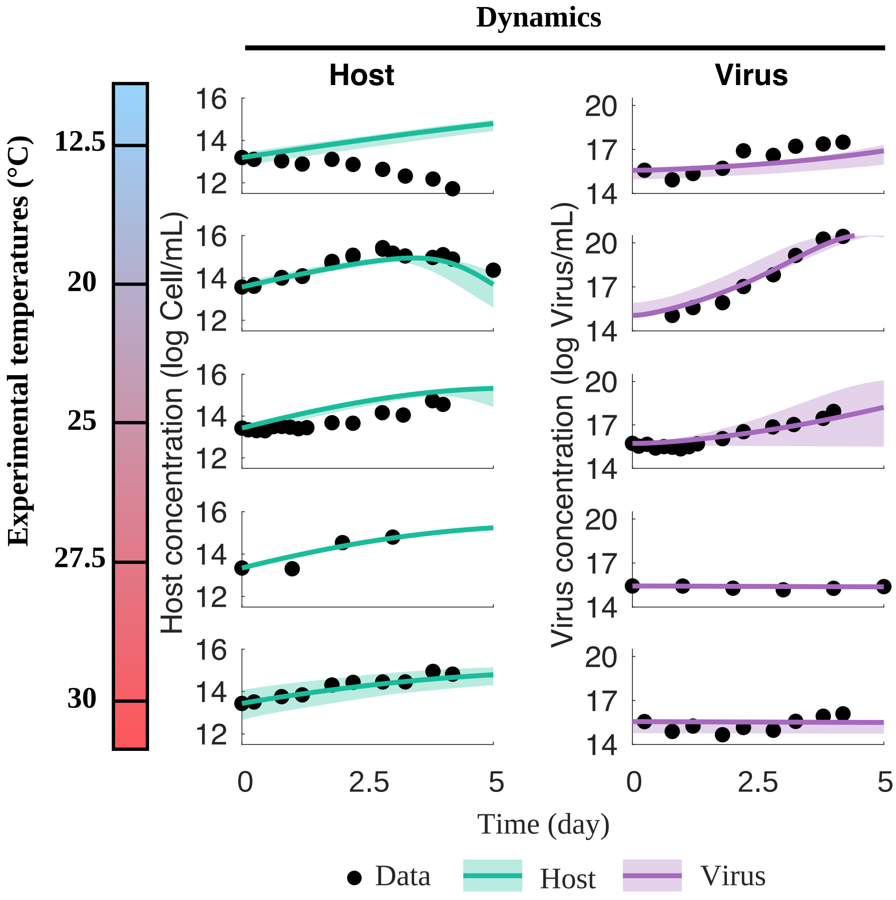


**Fig. S 2.** Model fits for the 5 temperatures tested experimentally in (1) for the pairs Mic-C/MicV-C (RCC834/RCC4229). The left column shows the phytoplankton dynamics with total host population $N$ model (green line) and experimental data (black circles). The right column shows the virus dynamics with total virus population $V$ model (purple line) experimental data (black circles). Shaded area represent the model fits variability when accounting for a $\pm$ 1°C uncertainty from experimental temperature.

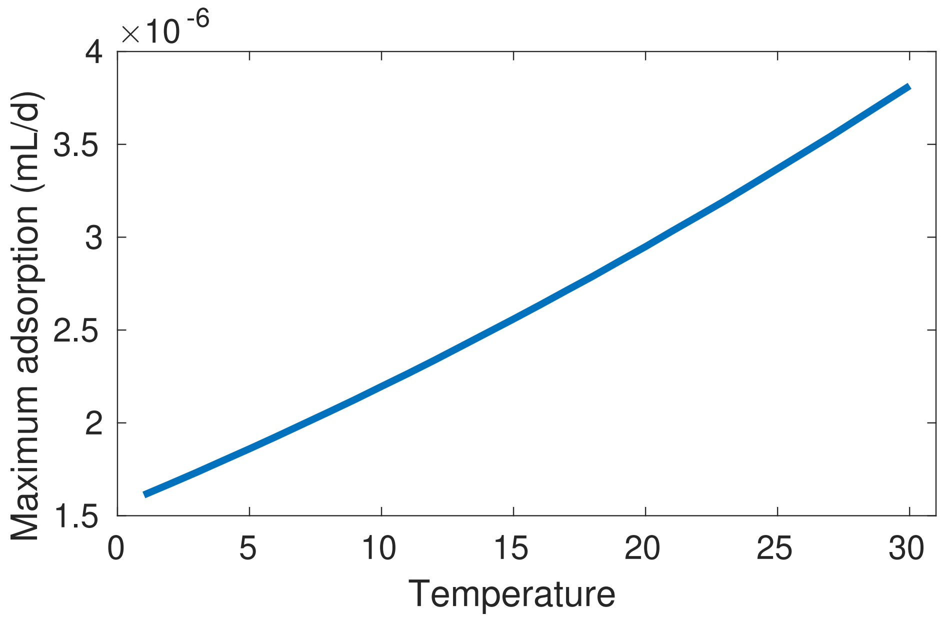


**Fig. S 3.** Biophysical maximum adsorption rate calculated according to (2; 3). We considered the diameter of MicV particles and *Micromonas* cells to be 130 nm and 1.5 $\mu$m respectively.

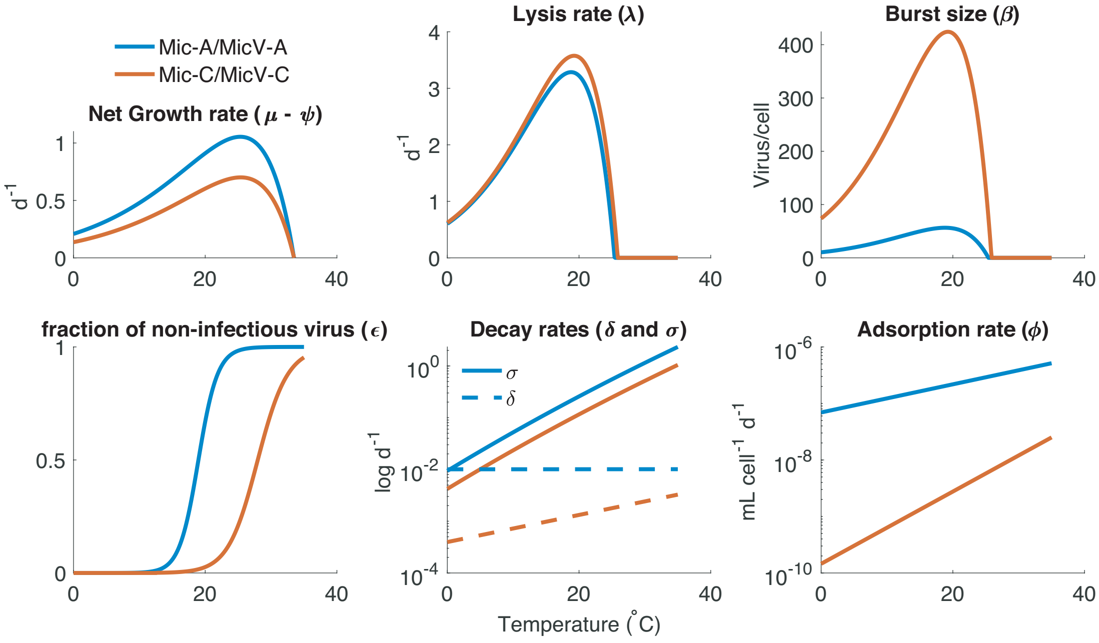


**Fig. S 4.** Life history trait temperature function for system Mic-A/MicV-A (blue) and Mic-C/MicV-C (orange).

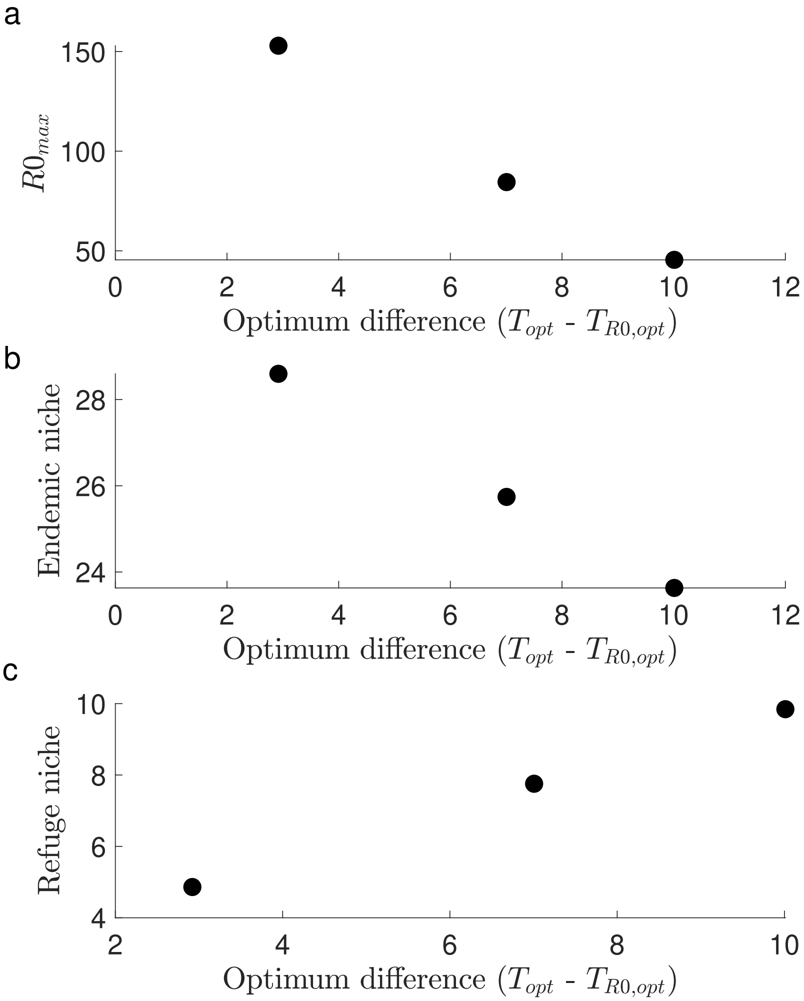


**Fig. S 5.** Relationships between optimum temperature difference and (a) the maximum $\mathscr{R}_0$, (b) the endemic niche ($T_{\mathscr{R}_0=1}$) and (c) the refuge niche ($T_{max} - T_{\mathscr{R}_0=1}$).

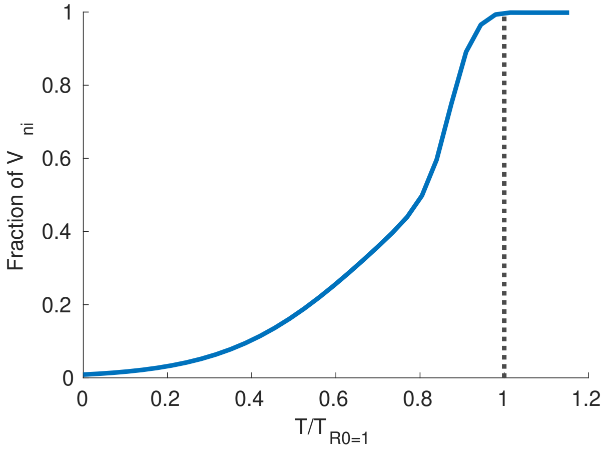


**Fig. S 6.** Fraction of non-infectious virus ($V_{ni}$) as a function of the ratio $T/T_{\mathscr{R}_0=1}$. The dashed black line represents $T = T_{\mathscr{R}_0=1}$.

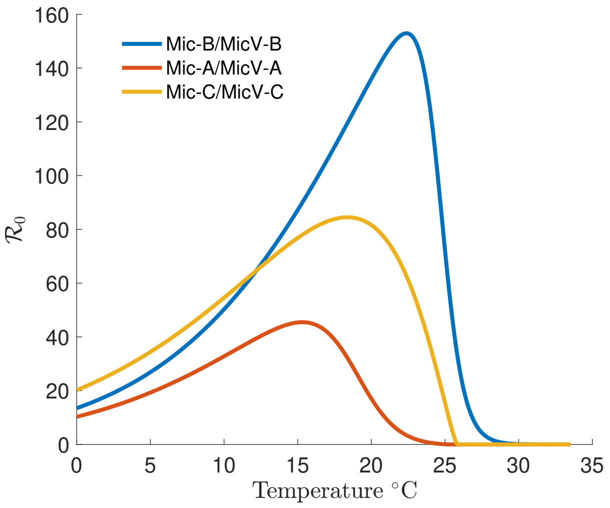


**Fig. S 7.** $\mathscr{R}_0$ as a function of temperature for the 3 virus-host pairs tested.

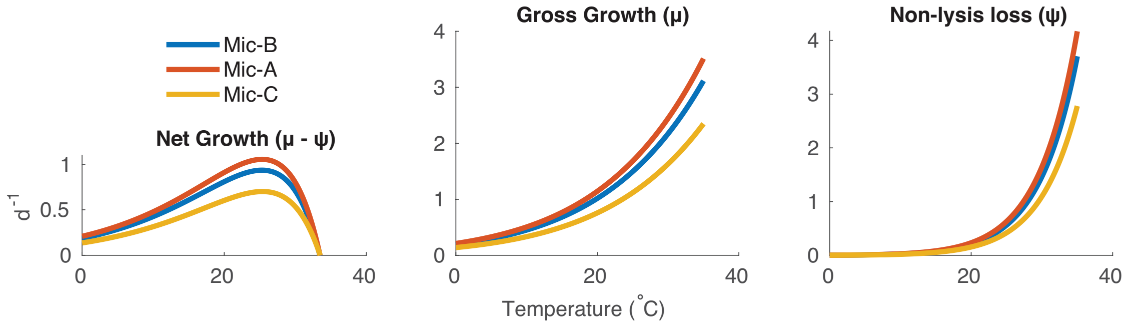


**Fig. S 8.** Growth and non-lysis mortality of the hosts Mic-B, Mic-A and Mic-C.

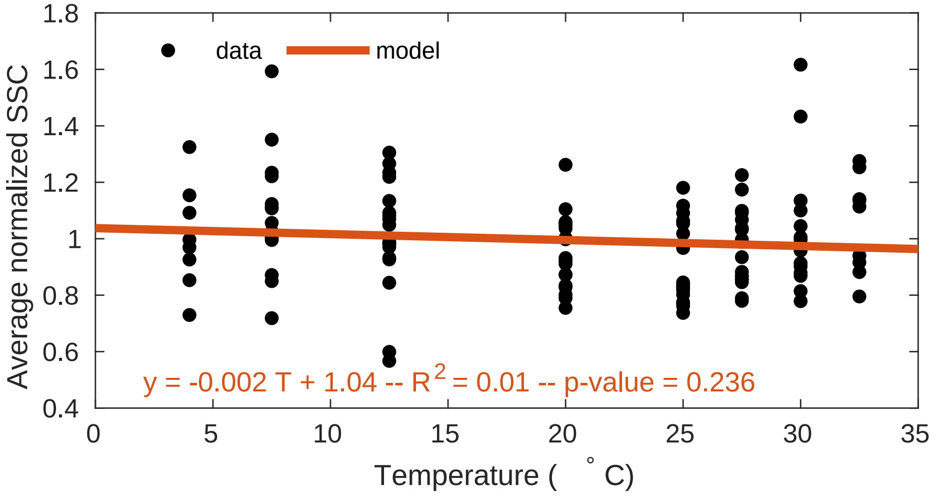


**Fig. S 9.** Side scatter (proxy of cell diameters) as function of temperature obtain by flow cytometry. Data are unpublished from (4) for 11 *Micromonas* strains. SSC was normalized by bead SSC.

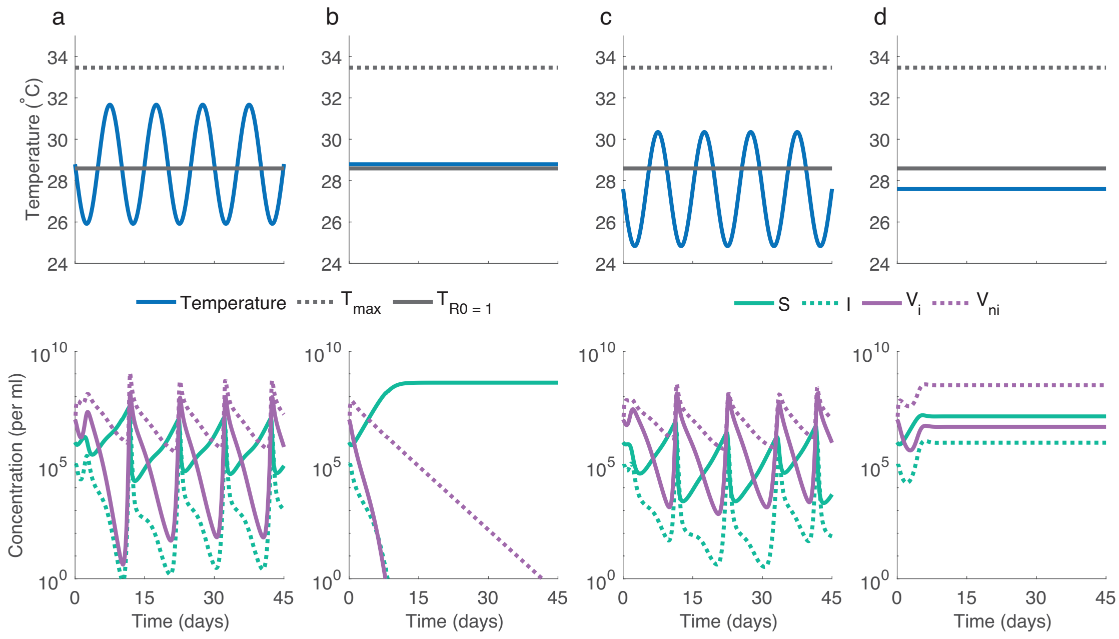


**Fig. S 10.** Impact of temperature on long-term virus-host dynamics for the pair Mic-B/MicV-B. The model was simulated for temperature $T_0$ beyond (a,b) and below (c,d) the temperature $T_{\mathscr{R}_0=1}$, corresponding to $\mathscr{R}_0 = 1$ for a constant (b,d) or a fluctuating (a,c) temperature during 45 days.

### References


**1.** Demory, D. *et al.* Temperature is a key factor in micromonas–virus interactions. *The ISME journal* **11**, 601 (2017).

**2.** Murray, A. G. & Jackson, G. A. Viral dynamics: a model of the effects of size, shape, motion and abundance of single-celled planktonic organisms and other particles. *Mar. Ecol. Prog. Ser.* 103–116 (1992).

**3.** Talmy, D. *et al.* Contrasting controls on microzooplankton grazing and viral infection of microbial prey. *Front. Mar. Sci.* **6**, 182 (2019).

**4.** Demory, D. *et al.* Picoeukaryotes of the micromonas genus: sentinels of a warming ocean. *The ISME journal* (2018).
